# Hierarchical temporal prediction captures motion processing from retina to higher visual cortex

**DOI:** 10.1101/575464

**Authors:** Yosef Singer, Ben D. B. Willmore, Andrew J. King, Nicol S. Harper

**Affiliations:** Department of Physiology, Anatomy and Genetics, University of Oxford, Sherrington Building, Parks Road, Oxford OX1 3PT, United Kingdom

## Abstract

Visual neurons respond selectively to specific features that become increasingly complex in their form and dynamics from the eyes to the cortex. Retinal neurons prefer localized flashing spots of light, primary visual cortical (V1) neurons moving bars, and those in higher cortical areas, such as middle temporal (MT) cortex, favor complex features like moving textures. Whether there are general computational principles behind this diversity of response properties remains unclear. To date, no single normative model has been able to account for the hierarchy of tuning to dynamic inputs along the visual pathway. Here we show that hierarchical application of temporal prediction - representing features that efficiently predict future sensory input from past sensory input - can explain how neuronal tuning properties, particularly those relating to motion, change from retina to higher visual cortex. This suggests that the brain may not have evolved to efficiently represent all incoming information, as implied by some leading theories. Instead, the selective representation of sensory inputs that help in predicting the future may be a general neural coding principle, which when applied hierarchically extracts temporally-structured features that depend on increasingly high-level statistics of the sensory input.

## Introduction

The temporal prediction [1] framework posits that sensory systems are optimized to represent features in natural stimuli that enable prediction of future sensory input. This would be useful for guiding future action, uncovering underlying variables, and discarding irrelevant information [1,2].Temporal prediction relates to a class of principles, such as the predictive information bottleneck [2–4] and slow feature analysis [5], that similarly involve selectively encoding only features that are efficiently predictive of the future. This class of principles differs from others that are more typically used to explain sensory coding – efficient coding [6,7], sparse coding [8] and predictive coding [9,10] – that aim instead to efficiently represent all current and perhaps past input. Although these principles have been successful in accounting for various visual receptive field (RF) properties in V1 [1,3,5,7–9], no single principle has so far been able to explain the diverse spatiotemporal tuning that emerges along the dorsal visual stream, which is responsible for the processing of object motion.

Any general principle of visual encoding needs to explain temporal aspects of neural tuning – the encoding of visual scenes in motion rather than static images. It is also important that any general principle is largely unsupervised. Some features of the visual system have been reproduced by deep supervised network models optimized for image classification using large labelled datasets (e.g. images labelled as cat, dog, car) [11]. While these models can help to explain the RF properties of the likely hard-wired retina [12], they are less informative if neuronal tuning is influenced by experience, as in cortex, since most sensory input is unlabeled except for sporadic reinforcement signals. The temporal prediction approach is unsupervised (i.e., it requires no labelled data), and inherently applies to the temporal domain. Furthermore, we have previously shown that a simple non-hierarchical model instantiating temporal prediction can account for temporal aspects of V1 simple cell RFs [1]. However, it is not known whether hierarchical application of the temporal prediction principle – an essential requirement for comparison with the organization of sensory pathways in the brain – can account for the emergence of motion processing along the visual pathway, from retina to higher visual cortex.

Here we have developed a hierarchical form of the temporal prediction model that predicts increasingly high-level statistics of natural dynamic visual input. It accounts not only for linear tuning properties of V1 simple cells, as in previous non-hierarchical temporal prediction models [1,3], but also for the diversity of linear and non-linear spatiotemporal tuning properties that emerge along the visual pathway. In particular, the temporal tuning properties in successive hierarchical stages of the model progress from those that distinguish magnocellular and parvocellular neurons at early levels of visual processing, to direction-selective simple and complex cells in V1, and finally to units that are sensitive to two-dimensional features of motion, as seen in end-stopping and pattern-selective cells in the cortex. The capacity of this model to explain tuning properties at multiple levels of the visual motion pathway using iterated application of a single process, suggests that optimization for temporal prediction may be a fundamental principle of sensory neural processing.

## Results

### The hierarchical temporal prediction model

We instantiated temporal prediction as a hierarchical model consisting of stacked feedforward single-hidden-layer convolutional neural networks (Fig. 1). The first stack was trained to predict the immediate future frame (40 ms) of unfiltered natural video inputs from the previous 5 frames (200 ms). Each subsequent stack was then trained to predict the future hidden-unit activity of the stack below it from the past activity in response to the natural video inputs. The four stacks contained 50, 100, 200 and 400 hidden units, respectively. L1 regularization was applied to the weights of each stack, akin to a constraint on neural wiring.

**Figure 1.**
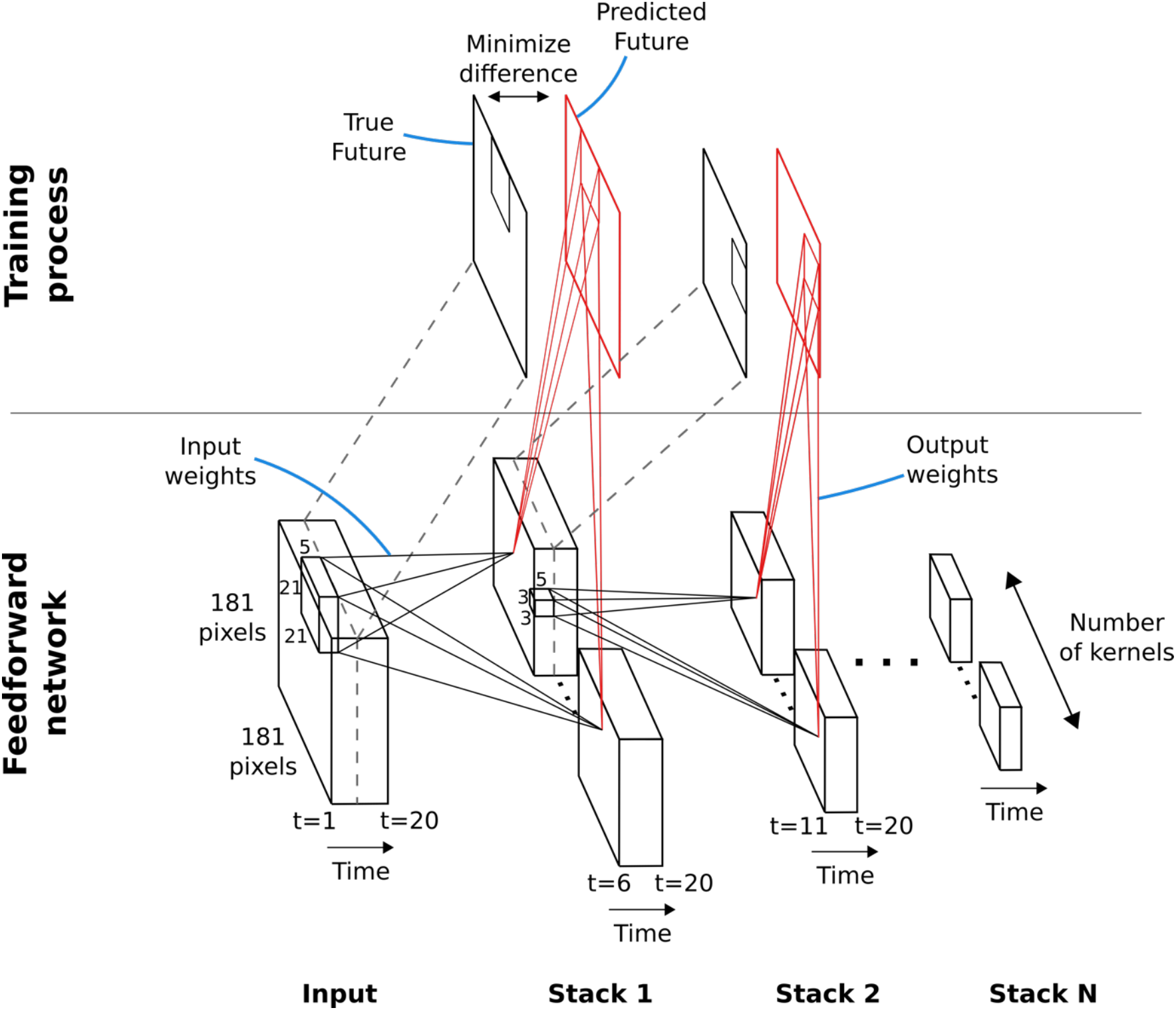
Hierarchical temporal prediction model. Schematic of model architecture. Each stack is a single hidden-layer feedforward convolutional network, which is trained to predict the future time-step of its input from the previous 5 time-steps. The first stack is trained to predict future pixels of natural video inputs from their past. Subsequent stacks are trained to predict future time-steps of the hidden-layer activity in the stack below, based on their past responses to the same natural video inputs.

### Temporal prediction of natural inputs produces retinal-like units

After training, we examined the input weights of the units in the first stack. Each hidden unit can be viewed as a linear non-linear (LN) model [13,14], as commonly used to describe neuronal RFs. With L1 regularization slightly above the optimum for prediction, the RFs of the units showed spatially-localized center-surround tuning with a decaying temporal envelope, characteristic of retinal and lateral geniculate nucleus (LGN) neurons [15–17]. The model units’ RFs have either an excitatory (ON) or inhibitory (OFF) blob-like structure at the 0ms time-step, often with a surround of opposing sign in the same or previous time-step (Figure 2a). Both ON and OFF units can have either small RFs that do not change polarity (Fig. 2a, units 1-4; Fig. 2b, bottom left) over time, or large RFs that switch polarity over time (Fig. 2a, units 5-6; Fig. 2b, top right). This is reminiscent of the four main cell types in the primate retina and LGN: the parvocellular-pathway ON/OFF neurons and the more change-sensitive magnocellular-pathway ON/OFF neurons, respectively [17].

**Figure 2.**
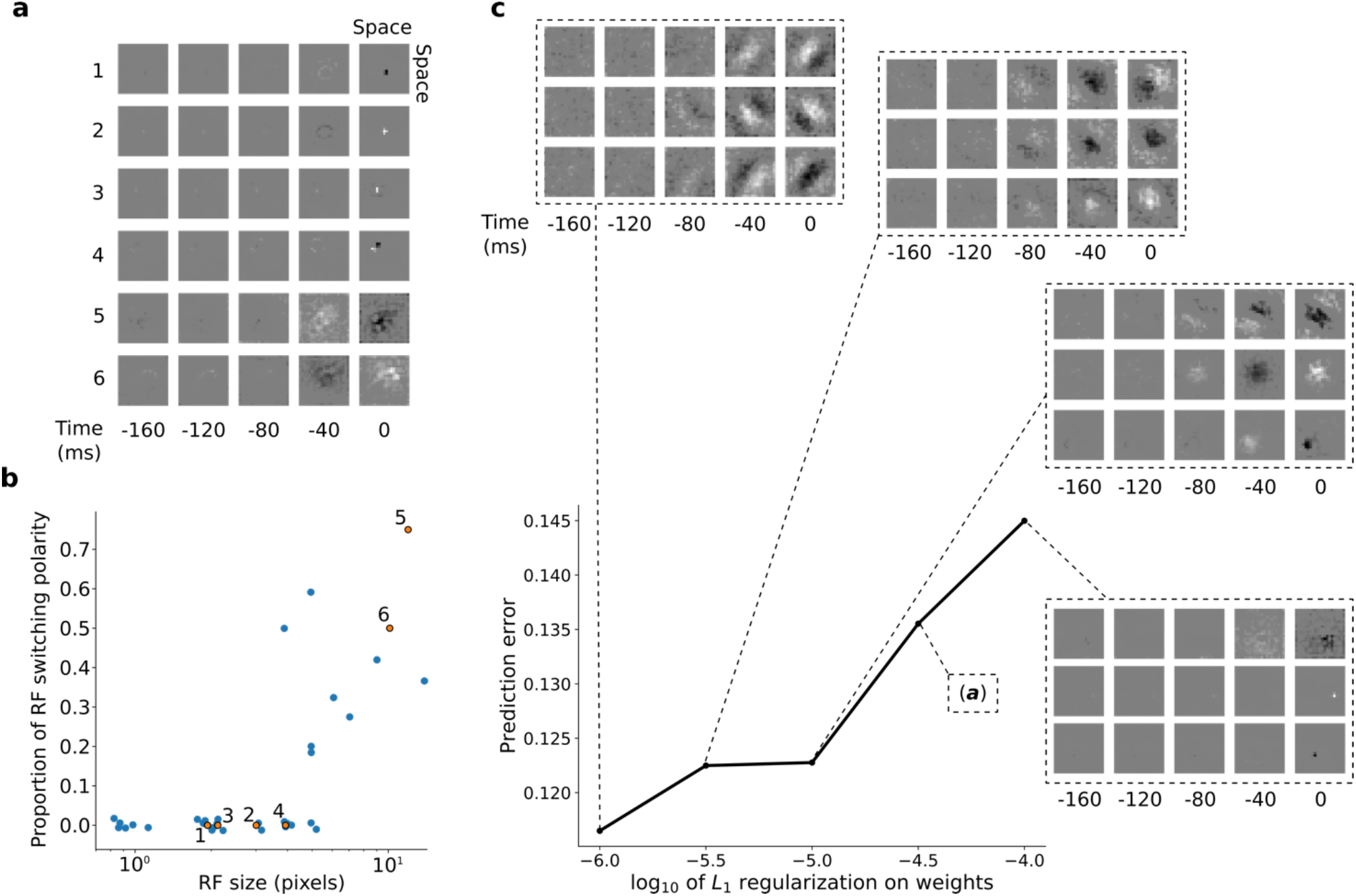
RFs of trained first stack of the model show retina-like tuning. **a,** Example RFs with center-surround tuning characteristic of neurons in retina and LGN. RFs are small and do not switch polarity over time (units 1-4) or large and switch polarity (units 5-6), resembling cells along the parvocellular and magnocellular pathways, respectively. **b,** RF size plotted against proportion of the pixels in the RF that switch polarity over the course of the most recent two timesteps. Units in *a* labelled and shown in orange. **c,** Effect of changing regularization strength on the qualitative properties of RFs.

Interestingly, simply decreasing L1-regularization strength causes the model RFs to change from center-surround tuning to Gabor-like tuning, resembling localized oriented bars that shift over time (Fig. 2c). It is possible that this balance, between a code that is optimal for prediction and one that prioritizes efficient wiring, might underlie differences in the retina and LGN of different species. The retina of mice and rabbits contains many neurons with oriented and direction-tuned RFs, whereas cats and macaques mostly have center-surround RFs [18]. Efficient retinal wiring may be more important in some species, due, for example, to different constraints on the width of the optic nerve or different impacts of light scattering by superficial retinal cell layers.

### Hierarchical temporal prediction produces simple and complex cell tuning

Using the trained center-surround-tuned network as the first stack, a second stack was added to the model and trained. The output of each second stack unit results from a linear-nonlinear-linear-nonlinear transformation of the visual input, and hence we estimated their RFs by reverse correlation with binary noise input. The resulting RFs were Gabor-like over space, resembling those of V1 simple cells [19–21]. The RF envelopes decayed into the past, and often showed spatial shifts or polarity changes over time, indicating direction or flicker sensitivity, as is also seen in V1 [22] (Fig. 3a,b, I; Fig. S1). Using full-field drifting sinusoidal gratings (Fig. 3a,b II), we found that most units were selective for stimulus orientation, spatial and temporal frequency (Fig. 3a,b, IV-VI), and some were also direction selective (Fig. 3b). Responses to the optimum grating typically oscillate over time between a maximum when the grating is in phase with the RF and 0 when the grating is out of phase (Fig. 3a,b, III). These response characteristics are typical of V1 simple cells [23].

**Figure 3.**
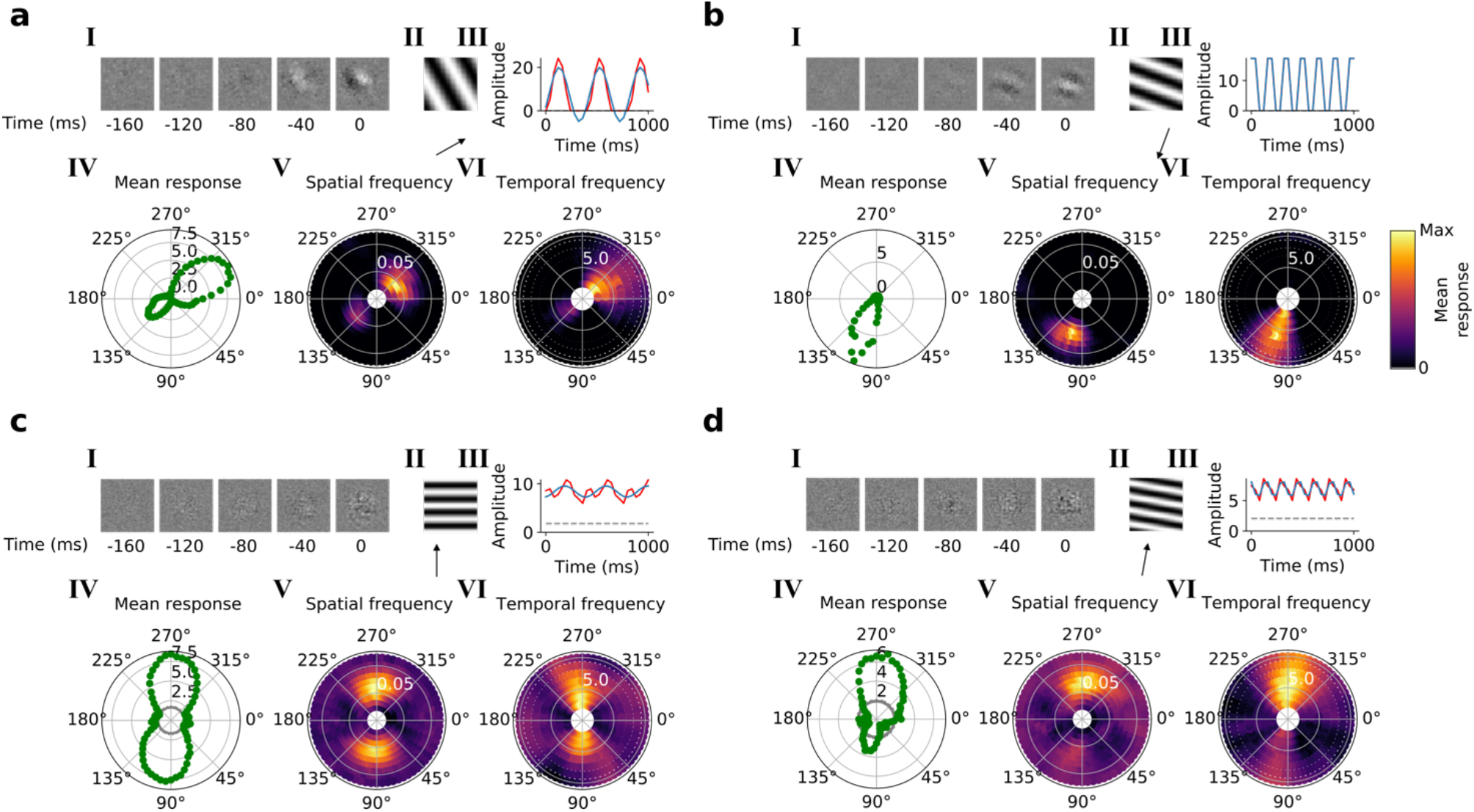
Qualitative tuning properties of model units in stacks 2 and 3. **a,b**, Tuning properties of two example units from the 2^nd^ stack of the model, including (I) the linear RF, (II) the drifting grating that best stimulates this unit and (III) the amplitude of the unit’s response to this grating over time. (IV) The unit’s mean response over time plotted against orientation (in degrees) for gratings presented at its optimal spatial and temporal frequency. (V, VI) Tuning curves showing the joint distribution of responses to (V) orientation (in degrees) and spatial frequency (in cycles/pixel) at the preferred temporal frequency and to (VI) orientation and temporal frequency (in Hz) at the preferred spatial frequency. In V and VI the color represents the mean response over time to the grating presented. **c,d**, As in (a,b) for example units in the 3^rd^ stack. Red line: unit response; blue line: best-fitting sinusoid; gray dashed line: response to blank stimulus. See also Figs. S1–S3.

In the third and fourth stack, we followed the same procedures as in the second stack. Most of these units are also tuned for orientation, spatial frequency, temporal frequency and in some cases for direction (Fig. 3c,d, IV-VI; Figs. S2, S3). However, while some units resembled simple cells, most resembled the complex cells of V1 and secondary visual areas (V2/V3) [21]. Complex cells are tuned for orientation and spatial and temporal frequency, but are relatively invariant to the phase of the optimum grating [24]; each cell’s response to its optimum grating has a high mean value and changes little with the grating’s phase (Fig. 3c,d, II, III). Whether a neuron is assigned as simple or complex is typically based on the modulation ratio in such plots (<1 indicates complex) [25]. Model units with low modulation ratios had little discernible structure in their RFs (Fig. 3c,d, I), another characteristic feature of V1 complex cells [26,27].

We quantified the tuning characteristics of units in stacks 2-4 and compared them to published V1 data [28] (Fig. 4a-j). The distribution of modulation ratios is bimodal in both V1 [25,28] and our model (Fig. 4a). Both model and real neurons were typically orientation selective, but with the model units having weaker tuning as measured by orientation bandwidth (median data [28]: 23.5°, model: 37.5°; Fig 4b) and circular variance (median data [28]: simple cells 0.45, complex cells 0.66; median model: simple cells 0.45, complex cells 0.85; Fig 4c,d). Orientation-tuned units (circular variance < 0.9) in the second stack were exclusively simple (modulation ratios > 1), whereas those in subsequent stacks became increasingly complex (Fig. 4a,c-f). In both model and data, circular variance was inversely correlated with the modulation ratio (Fig. 4c-e,h). Similarly, orientation bandwidth varied with modulation ratio in a similar way in both model and data (Fig. 4f,i). Model units showed a range of direction selectivity preferences (Fig. 4g), with simple cell-like units (69% with direction selectivity index, DSI ≥ 0.5; n=155; Fig. 4j) tending to be more direction tuned than complex cell-like units (20% with DSI ≥ 0.5, n=205; Fig. 4j) as is seen in V1 [29].

**Figure 4.**
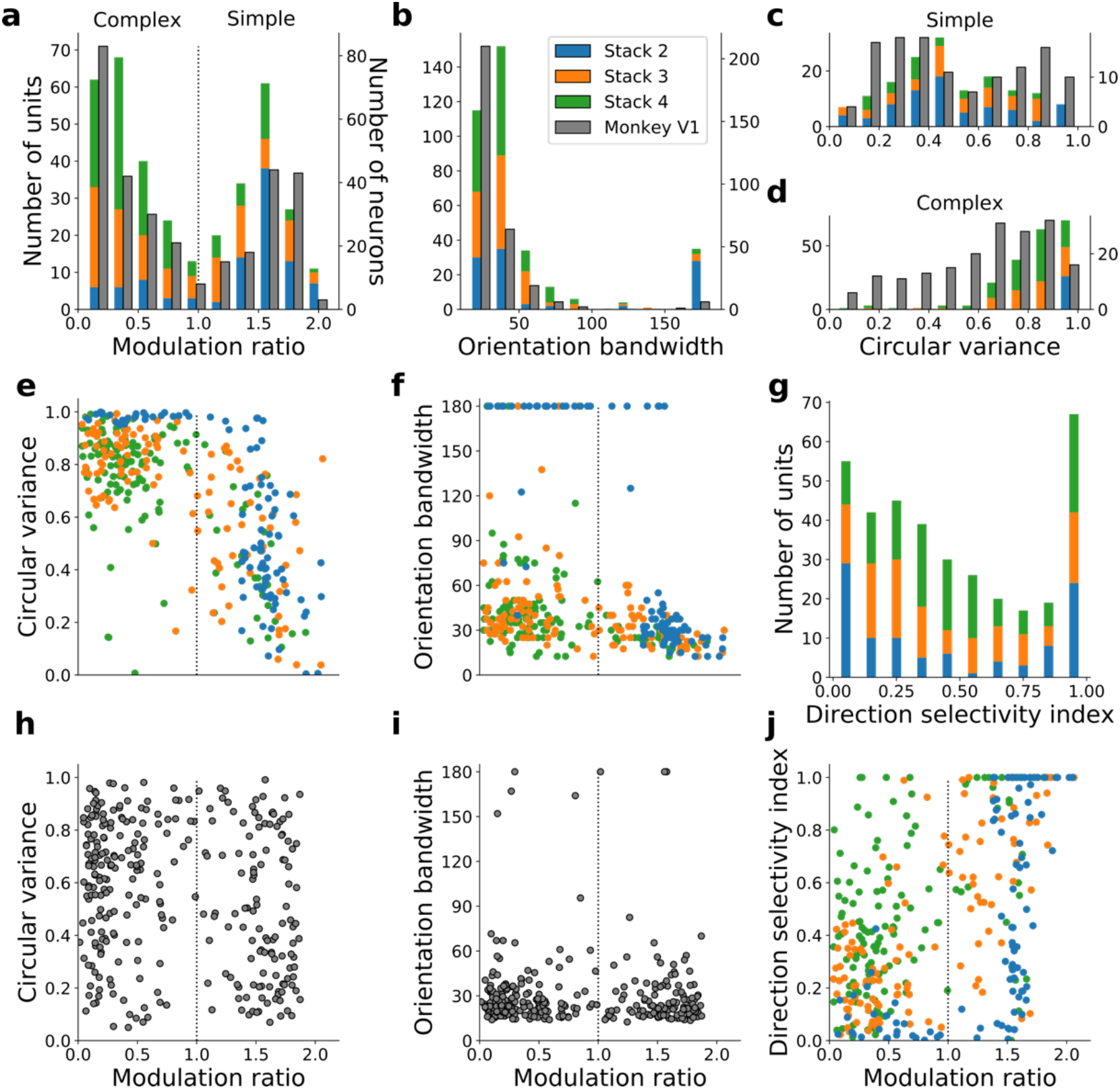
Quantitative tuning properties of model units in stacks 2-4 in response to drifting sinusoidal gratings and corresponding measures of macaque V1 neurons. **a-d,** Histograms showing tuning properties of model and macaque V1 neurons as measured using drifting gratings. Units from stacks 2-4 are plotted on top of each other to also show the distribution over all stacks; this is because no stack can be uniquely assigned to V1. Modulation ratio measures how simple or complex a cell is; orientation bandwidth and circular variance are measures of orientation selectivity. **e, f,** Joint distributions of tuning measures for model units. **g,** Distribution of direct selectivity for model units. **h, i,** Joint distributions of tuning measures for V1 data; note similarity to e and f. **j,** Joint distribution of direction selectivity and modulation ratio for model units.

### Model units are tuned to two-dimensional features of visual motion

Simple and complex cells extract many dynamic features from natural scenes. However, their small RFs prevent individual neurons from tracking the motion of objects because of the aperture problem; the direction of motion of an edge is ambiguous, with only the component of motion perpendicular to the cell’s preferred orientation being represented. Two classes of neurons exist that can recover 2-dimensional motion information and overcome the aperture problem. End-stopped neurons, found in primary and secondary visual areas, respond unambiguously to the direction of motion of endpoints of restricted moving contours [30]. Pattern-selective neurons, in MT of primates, solve the problem for the more general case, likely by integrating over input from many direction-selective V1 complex cells [31–35], and hence respond selectively to the motion of patterns as a whole.

To investigate end-stopping in our model units, we applied circular masks of different sizes to the grating stimuli. Some units displayed end-stopping, responding most strongly to gratings with an intermediate mask radius, with the response decreasing as the radius increased beyond this (Fig. 5a,b). To determine whether these end-stopped units unambiguously represent the direction of motion of end-points, we measured two-bar response maps [30], which determine response dependence on the horizontal and vertical components of motion (see Methods). Recordings from V1 indicate that more strongly end-stopped neurons have a weak tendency for less ambiguous motion tuning in these maps [30] (i.e. have less elongated excitatory regions). Consistent with this, our model produces examples of end-stopped units with less ambiguous motion tuning (Fig. 5b,c, first two panels) and non-end-stopped units with more ambiguous motion tuning (Fig. 5b,c, last panel).

**Figure 5.**
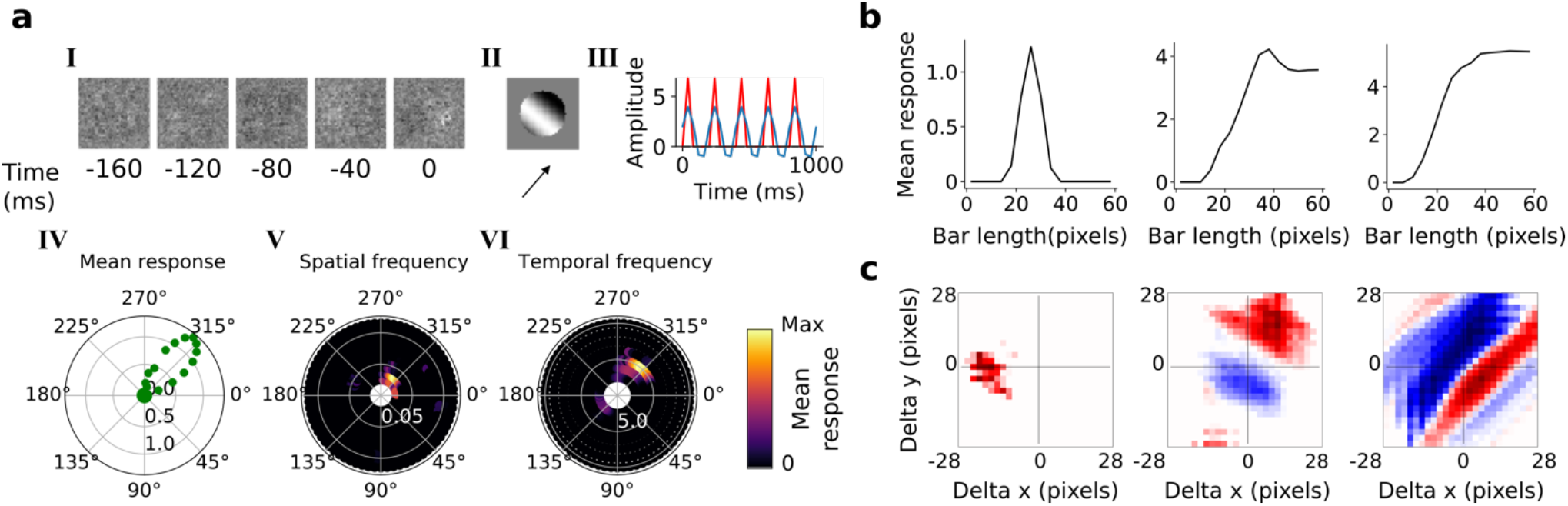
End-stopping. **a**, Example end-stopped model unit. I-VI as in Figure 3. **b**, Response as a function of bar length for the unit in **a** (left) and two other example units (middle, right). **c**, 2-bar maps of units with corresponding bar-length tuning plots shown in **b**.

To investigate pattern selectivity in our model units, we measured their responses to drifting plaids, comprising two superimposed drifting sinusoidal gratings with different orientations. The net direction of the plaid movement lies midway between these two orientations (Fig. 6a). In V1 and MT, component-selective cells respond maximally when the plaid is oriented such that either one of its component gratings moves in the preferred direction of the cell (as measured by a drifting grating). This results in two peaks in plaid-direction tuning curves [31,34,36]. Conversely, pattern-selective cells (in macaque typically seen in MT and not V1) have a single peak in their direction tuning curves, when the plaid’s direction of movement aligns with the preferred direction of the cell [31,34,36]. We see examples of both component-selective units (in stacks 2-4) and pattern-selective units (only in stack 4) in our model, as indicated by plaid-direction tuning curves (Fig. 6b) and plots of response as a function of the directions of each component (Fig. 6c).

**Figure 6.**
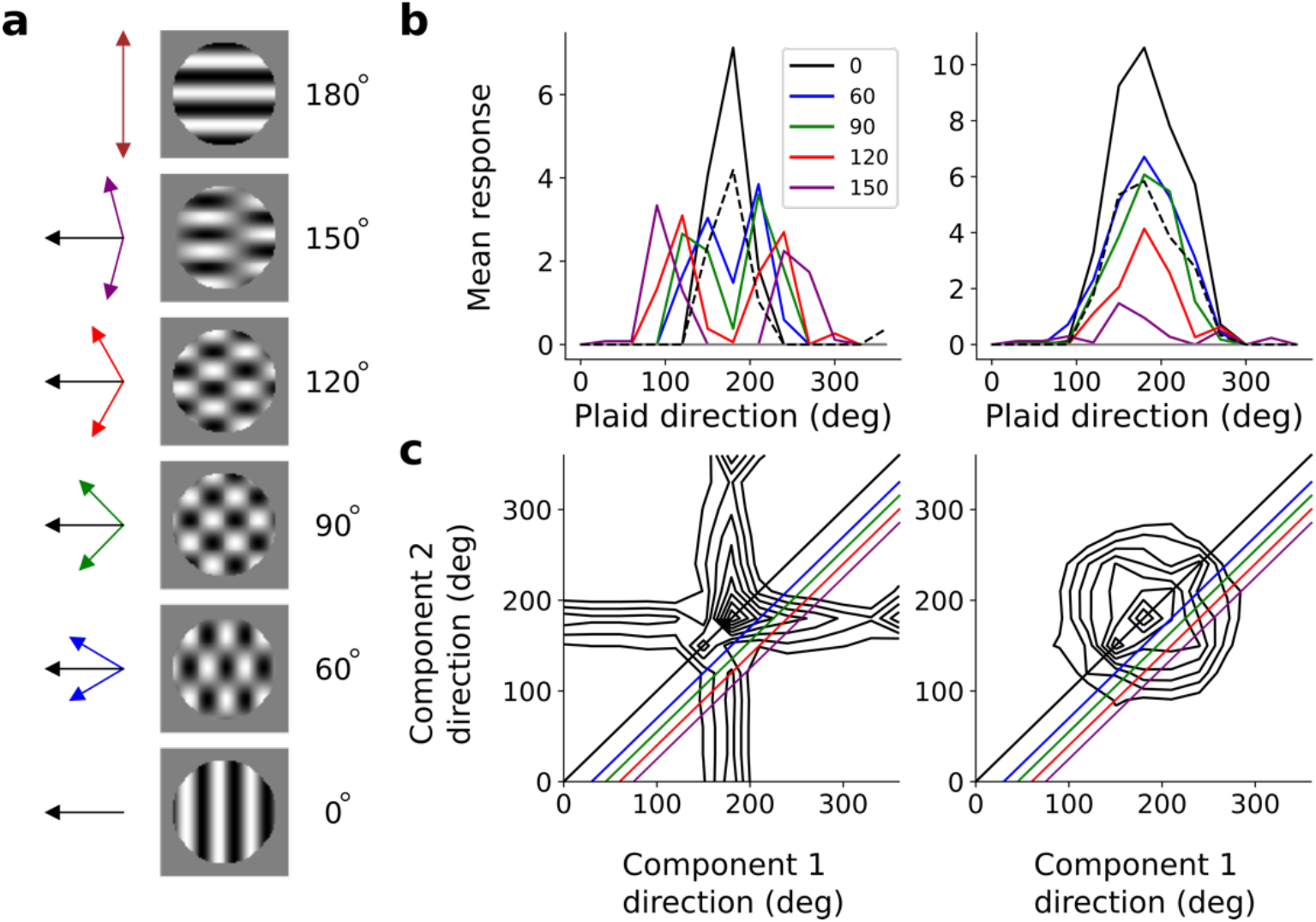
Pattern sensitivity. **a**, Example plaid stimuli used to measure pattern selectivity. Black arrow, direction of pattern motion. Colored arrows, directions of component motion. **b**, Direction tuning curves showing the response of an example component-selective (left) and pattern-selective (right) unit to grating and plaid stimuli. Colored lines, response to plaid stimuli composed of gratings with the indicated angle between them. Black solid line, unit’s response to double intensity grating moving in the same direction as plaids. Dotted line, response to single intensity grating moving in the same direction. Gray line, response to blank stimulus. **c**, Surface contour plots showing response of units in **b** to plaids as a function of the direction of the grating components. Colored lines denote loci of plaids whose responses are shown in the same colors in **b**. Contour lines range from 20% of the maximum response to the maximum in steps of 10%. For clarity, all direction tuning curves are rotated so that the preferred direction of the response to the optimal grating is at 180°. Responses are mean amplitudes over time.

## Discussion

We have presented a simple model that hierarchically instantiates temporal prediction – the hypothesis that sensory neurons are optimized to efficiently predict the immediate future from the recent past. This model has several advantages: it is unsupervised; it operates over spatiotemporal inputs and its hierarchical implementation is general, allowing the same network model to learn features resembling each level of the visual processing hierarchy with little fine-tuning or modification to the model structure at each stage. This simple model accounts for many spatial and temporal tuning properties of cells in the dorsal visual pathway, from the center-surround tuning of cells in the retina and LGN, to the spatiotemporal tuning and direction selectivity of V1 simple and complex cells, and the motion processing of end-stopped and pattern selective cells.

Although this work suggests that temporal prediction may explain why many features of sensory neurons take the form that they do, we are agnostic as to whether the features are hard-wired by evolution or learned over the course of development. This is likely to depend on the region in question, with retina more hard wired and cortex a mixture of innate tuning and learning from sensory input [37,38]. If they are learned, this suggests that while some neurons represent the predictive features, a fraction of neurons in the cortex (or elsewhere) might represent the prediction error used to train the network. There is evidence that cortical neurons might represent prediction error [39,40]. Our model is trained using backpropagation over a single hidden layer for each stack. Although the biological plausibility of backpropagation has been questioned, increasingly biologically realistic methods for training networks are being developed [41]. There are a number of further developments that could be made to our model that may even better capture features of the biology, notably the inclusion of recurrent connections within and between layers or using spiking units. Furthermore, although we propose that temporal prediction is an important method for extracting potentially useful features of natural inputs, additional constraints are likely to further refine those features that are of direct relevance to specific behavioral goals.

There are other normative models of visual processing, based on a range of principles, which can account for a number of properties of visual neurons. Prominent theories include predictive coding, sparse coding, independent component analysis (ICA) and temporal coherence. The predictive coding framework postulates that sensory systems learn the statistical regularities present in natural inputs, feeding forward the errors caused by deviations from these regularities to higher areas [9,10,42]. In this process, the predictable components of the input signal are removed and only unexpected inputs are fed forward through the hierarchy. This should be distinguished from temporal prediction, which performs selective coding, where predictive elements of the input are explicitly represented in neuronal responses and non-predictable elements are discarded [2–4]. Sparse coding [8,43], which shares similarities with predictive coding [10], is built on the idea that an overcomplete set of neurons is optimized to represent inputs as accurately as possible using only few active neurons for a given input. ICA [7,44] is a related framework that finds maximally independent features of the inputs. Sparse coding and ICA are practically identical in cases where a critically complete code is used. In these frameworks, as with predictive coding, the aim is to encode all current or past input, whereas in temporal prediction, only features that are predictive of the future are encoded and other features are discarded.

Another set of approaches, slow feature analysis [45] (SFA) and slow subspace analysis [46] (SSA), stem from the idea of temporal coherence [47], which suggests that a key goal of sensory processing is to identify slowly varying features of natural inputs. SFA is closely related to temporal prediction because features that vary slowly are likely to be predictive of the future. However, SFA and temporal prediction may give different weighting to the features that they find [48], and SFA could also fail to capture features that do not vary slowly, but are predictive of the future.

We will focus on unsupervised normative models (i.e. those trained on natural inputs) because they are the most relevant to our model. Broadly, these models can be divided into several categories: local models, trained to represent features of a specific subset of neurons, such as simple cells in V1, and hierarchical models, which attempt to explain features of more than one cell type (such as simple and complex cells) in a single model. These two categories can be further divided into models that are trained on natural spatial inputs (images) and those that are trained on natural spatiotemporal inputs (movies).

Among local models, sparse coding and ICA are the standard normative models of V1 simple cell RFs [7,8,43,44,49,50]. Typically, these models are trained using still natural images and have shown remarkable success in accounting for the spatial features of V1 simple cell RFs [7,8,43,44]. However, models trained on static images are unable to account for temporal aspects of neuronal RFs, such as direction selectivity or two-dimensional motion processing. The ICA and sparse coding frameworks have been extended to model features of spatiotemporal inputs [3,49,50]. While these models capture many of the spatial tuning properties of simple cells, they tend to produce symmetric temporal envelopes that do not match the asymmetric envelopes of real neurons [1]. Capturing temporal features is especially important when building a normative model of the dorsal visual stream, which is responsible for processing cues related to visual motion. When trained to find slowly varying features in natural video inputs, SFA models [5] find features with tuning properties that closely resemble those of V1 complex cells, including phase invariance to drifting sinusoidal gratings and end- and side-inhibition. A sparse prior must be applied to the activities of the model units in order to produce spatial localization – a key feature of V1 complex cells [51]. Although SFA can account for complex cell tuning, on its own this framework does not provide a normative explanation for simple cells.

A notable hierarchical model, predictive coding [9] provides a powerful framework for learning hierarchical structure from visual inputs in an unsupervised learning paradigm. When applied to natural images, predictive coding has been used successfully to explain the oriented tuning of simple cells in V1 and some nonlinear tuning properties of neurons in this area, such as end-stopping [9]. However, it is not clear whether this framework can reproduce the phase-invariant tuning of complex cells. Nor has it been shown to account for direction selectivity, end-stopped tuning for motion [30], or pattern motion sensitivity [31].

Hierarchical ICA models (and related models) provide another approach. These consist of two-layer networks that are trained on natural images with an independence prior placed on the unit activities [52–55]. They have been shown to produce simple cell subunits in the first layer of the network and phase-invariant tuning reminiscent of complex cells in the second layer. However, these models typically incorporate aspects to specifically encourage complex cell-like characteristics in the form of a quadratic nonlinearity resembling the complex cell energy model [56]. In some models [57], phase invariance is enforced by requiring the outputs of individual subunits to be uncorrelated [58]. This is in contrast to our model where the phase invariance is learned as a consequence of the general principle of finding features that can efficiently predict future input. An advantage to learning the invariance rather than hand-crafting it is that the identical model architecture can then be applied hierarchically to explain features in higher visual areas without changing the form of the model.

Some hierarchical models based on temporal coherence have been trained on spatiotemporal inputs (videos of natural scenes). These models have been shown to capture properties of both simple and complex cells [46,58,59]. Typically, these share a similar structure and assumptions with the hierarchical ICA models outlined above, consisting of a two-layer model where the outputs of one or more simple cell subunits are squared and passed forward to a complex cell layer. Other models have combined sparsity/independence priors with a temporal slowness constraint in a hierarchical model [60–63]. Since sparsity constraints tend to produce simple cell tuning and slowness constraints result in complex cell tuning, these models produce units with both types of selectivity. This contrasts with our model which produces both types of selectivity with a single objective and also accounts for tuning in higher visual areas.

Previous studies using principles related to temporal prediction have produced non-hierarchical models of retina or primary cortex alone and demonstrated retina-like RFs [64], simple-cell like RFs [1,3], or RFs resembling those found in primary auditory cortex [1]. However, any general principle of neural representation should be extendable to a hierarchical form and not tailored to the region it is attempting to explain. Here we show that temporal prediction can indeed be made hierarchical and so reproduce the major motion-tuning properties that emerge along the dorsal visual pathway from retina to MT. This model captures not only linear tuning features such as those seen in center-surround retinal neurons and direction-selective simple cells, but also nonlinear features seen in complex cells, end-stopped cells and pattern-sensitive neurons. The capacity of our hierarchical temporal prediction model to account for so many tuning features at multiple levels of the visual system suggests that the same framework may well explain many more features of the brain than we have investigated. Furthermore, iterated application of temporal prediction, as performed by our model, could be used to make predictions about the tuning properties of neurons in brain pathways that are much less well understood than those of the visual system. Our results suggest that, by learning behaviorally-useful features from dynamic unlabeled data, temporal prediction may represent a fundamental coding principle in the brain.

## Methods

### Data used for model training and testing

#### Visual inputs

Videos (grayscale, without sound, sampled at 25 fps) of wildlife in natural settings were used to create visual stimuli for training the artificial neural network. The videos were obtained from http://www.arkive.org/species, contributed by: BBC Natural History Unit, http://www.gettyimages.co.uk/footage/bbcmotiongallery; BBC Natural History Unit & Discovery Communications Inc., http://www.bbcmotiongallery.com; Granada Wild, http://www.itnsource.com; Mark Deeble & Victoria Stone Flat Dog Productions Ltd., http://www.deeblestone.com; Getty Images, http://www.gettyimages.com; National Geographic Digital Motion, http://www.ngdigitalmotion.com. The longest dimension of each video frame was clipped to form a square image. Each frame was then down-sampled (using bilinear interpolation) over space, to provide 180 × 180 pixel frames. The video patches were cut into non-overlapping clips, each of 20 frames duration (800 ms). We used a training set of *N* = ~1305 clips from around 17 min of video, and a validation set of *N* = ~145 clips. Finally, each clip was normalized by subtracting the mean and dividing by the standard deviation of that clip.

### Hierarchical temporal prediction model

#### The model and cost function

The hierarchical temporal prediction model consisted of stacked feedforward single-hidden-layer 3D convolutional neural networks. Each stack consisted of an input layer, a convolutional hidden layer and a ‘transposed convolutional’ output layer. Each unit (convolutional kernel) in the hidden layer performed 3D convolution over its inputs (over time and 2D space; Figure 1) and its output was determined by passing the result of this operation through a rectified linear function. Following the hidden layer there was a ‘transposed convolutional’ output layer, which again performed convolution (and dilation for stride >1). Each stack was trained to minimize the difference between its output and its target. The target was the input at the immediate future time-step.

The first stack of the model was trained to predict the immediate future frame (40 ms) of unfiltered natural video inputs from the previous 5 frames (200 ms). Each subsequent stack was then trained to predict the immediate future hidden-unit activity of the stack below it from the past hidden-unit activity in response to the natural video inputs. This process was repeated until 4 stacks had been trained. The first stack used 50 hidden units and this number was doubled with each added stack, until we had 400 units in the 4^th^ stack.

More formally, each stack of model can be described by a network of the same form. The input to the network has *i* = 1 to to *I* input channels. For channel *i*, for clip *n*, the input **<u>U</u>**_*in*_ is a rank-3 tensor spanning time and 2D-space with *x* = 1 to *X* and *y* = 1 to *Y* spatial positions, and *t* = 1 to *T* time steps. Throughout the Methods, capital, bold and underlined variables are rank-3 tensors over time and 2D-space, otherwise variables are scalars. The first stack has only a single input channel (the grayscale input frames, *I* = 1). Each subsequent stack had as many input channels (*I*) as the number of hidden units (feature maps) in the previous stack.

The network has a single hidden layer of *j* = 1 to *J* convolutional kernels. For clip *n* and kernel *j*, the output of each kernel is a rank-3 tensor over time and 2D-space, **<u>H</u>**_*jn*_:

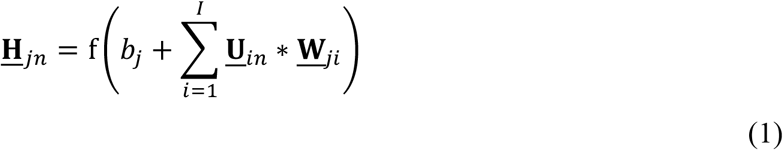

The parameters in Equation 1 are the connective input weights of kernels **<u>W</u>**_*ji*_ (between each input channel *i* and hidden unit *j*) and the bias *b*_*j*_ (of hidden unit *j*). f() is the rectified linear function and * is the 3D convolutional operator over the two spatial and one temporal dimensions of the input, with stride (s_1_, s_2_, s_3_). Each hidden layer kernel **<u>W</u>**_*ji*_ is 3D with size (*X’,Y’,T’*). No zero padding is applied to the input.

The output of the network predicts the future activity of the input. Hence, the number of input channels (*I*) always equals the number of output channels (*K*) for each stack. To ensure that the predicted output has the same size as the input when a stride of >1 is used, the hidden layer representation is dilated by adding *s*-1 zeros between adjacent input elements, where *s* = (s_1_, s_2_, s_3_) is the stride of the convolutional operator in the hidden layer. The dilated hidden-unit output is 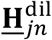. When stride=1, 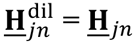.

The activity 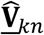 of each output channel *k* is the estimate of the true future **<u>V</u>**_*kn*_ given the past **<u>U</u>**_*in*_. **<u>V</u>**_*kn*_ is simply **<u>U</u>**_*in*_ shifted one time step into the future, and *k* = *i*. This prediction 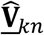 is estimated from the hidden unit output 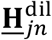 by:

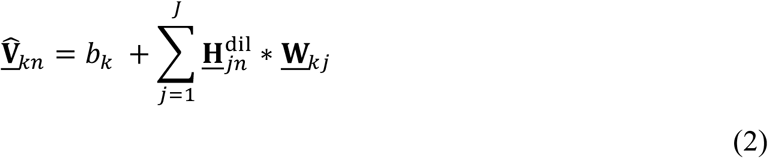

The parameters in Equation 2 are the connective output kernels **<u>W</u>**_*kj*_ (the weights between each hidden unit *j* and output channel *k*) and the bias *b*_*k*_. Each output kernel **<u>W</u>**_*kj*_ is 3D with size (*X’,Y’,*1), predicting a single time-step into the future based on hidden layer activity in that portion of space. The parameters **<u>W</u>**_*ji*_, **<u>W</u>**_*kj*_, *b*_*j*_, and *b*_*k*_ were optimized for the training set by minimizing the cost function given by:

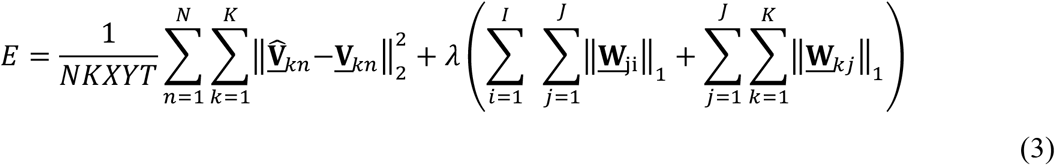

Where ‖ ‖_*p*_ is the entrywise *p*-norm of the tensor over time and 2D-space, p = 2 is the sqrt of the sum of squares of all values in the tensor, and p = 1 is the sum of absolute values. Thus, *E* is the sum of the squared error between the prediction 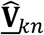 and the target **<u>V</u>**_*kn*_, plus an *L*_1_-norm regularization term, which is proportional to the sum of absolute values of all weights in the network and its strength is determined by the hyper-parameter *λ*. This regularization tends to drive redundant weights to near zero and provides a parsimonious network.

#### Implementation details

The networks were implemented in Python (https://lasagne.readthedocs.io/en/latest/; http://deeplearning.net/software/theano/). The objective function was minimized using backpropagation as performed by the Adam optimization method [65] with hyperparameters β1 and β2 kept at their default settings of 0.9 and 0.999, respectively, and the learning rate (α) varied as detailed below. Training examples were split into minibatches of 32 training examples each.

During model network training, several hyperparameters were varied, including the regularization strength (λ) and the learning rate (α). For each hyperparameter setting, the training algorithm was run for 1000 iterations. The effect of varying λ on the prediction error (the first term of Equation 3) and receptive field structure of the first stack is shown in Fig. 2. For all subsequent stacks, we varied λ between 10^−5^ and 10^−7^ and picked the network with the lowest prediction error (mean squared error) on a held-out validation set. We measured the predictive capacity of each network by taking the average prediction error on the validation set over the final 50 iterations. The settings for each stack are presented in Table 1:

**Table 1.** Model parameter settings for each stack.

| Stack | Input size<br>(X,Y,T,I) | Hidden layer<br>size | Kernel<br>size<br>(X',Y',T') | Spatial and<br>temporal<br>extent | Stride<br>(s<sub>1</sub>,s<sub>2</sub>,s<sub>3</sub>) | Number<br>of kernels | Learning<br>rate ($\alpha$) | L1<br>Regularization<br>strength ($\lambda$) |
| --- | --- | --- | --- | --- | --- | --- | --- | --- |
| 1 | 181x181x20x1 | 17x17x16x50 | 21x21x5 | 21x21x5 | 10x10x1 | 50 | $10^{-2}$ | $10^{-4.5}$ |
| 2 | 17x17x16x50 | 15x15x12x100 | 3x3x5 | 41x41x9 | 1x1x1 | 100 | $10^{-4}$ | $10^{-6}$ |
| 3 | 15x15x12x100 | 13x13x8x200 | 3x3x5 | 61x61x13 | 1x1x1 | 200 | $10^{-4}$ | $10^{-6}$ |
| 4 | 13x13x8x200 | 11x11x4x400 | 3x3x5 | 81x81x17 | 1x1x1 | 400 | $10^{-4}$ | $10^{-6}$ |

### Model unit spatiotemporal extent and receptive fields

Due to the convolutional form of the hidden layer, each hidden unit can potentially receive from a certain span over space and time. We call this the unit’s spatial and temporal extent. For stack 1, this this extent is given by the kernel size (21 × 21 × 5, space × space × time). For stack 2, the extent of each hidden unit is a function of its kernel size and the kernel size and stride of the hidden units in the previous stack, resulting in an extent of 41 × 41 × 9. Similarly, the extent of each hidden unit in stack 3 is 61 × 61 × 13 and in stack 4 is 81 × 81 × 17. The RF size of a unit can be considerably smaller than the hidden unit’s extent.

In the first stack of the model, the combination of linear weights and nonlinear activation function are similar to the basic linear non-linear (LN) model [13,14] commonly used to describe neuronal RFs. Hence, the input weights between the input layer and a hidden unit of the model network are taken directly to represent the unit’s RF, indicating the features of the input that are important to that unit. The output activities of hidden units in stacks 2-4 are transformations with multiple linear and nonlinear stages, and hence we estimated their RFs by applying reverse correlation to 100,000 responses to binary noise input with amplitude ±3 to stack 1.

### In vivo V1 receptive field data

Responses to drifting gratings measured using recordings from V1 simple and complex cells were compared against the model (Fig. 4). The *in vivo* data were taken from http://www.ringachlab.net/lab/Data.html [28].

### Receptive field size and polarity

We measured the size of the RFs of the units in the first stack and examined the relationship between the RF size and the proportion of the RFs switching polarity. For each unit, all pixels in the most recent time-step with intensities ≥50% of the maximum pixel intensity in that time-step are included in the RF. The RF size was determined by counting the number of pixels fitting this criterion. We then counted the proportion of pixels included in the RF that changed sign (either positive to negative or vice versa) between the two most recent timesteps. The relationship between these two properties for the units in the first stack is shown in Fig. 2b.

### Drifting sinusoidal gratings

In order to characterize the tuning properties of the model’s visual RFs, we measured the responses of each unit to full-field drifting sinusoidal gratings. For each unit, we measured the response to gratings with a wide range of orientations, spatial frequencies and temporal frequencies until we found the parameters that maximally stimulated that unit (giving rise to the highest mean response over time). We define this as the optimal grating for that unit. In cases where orientation or tuning curves were measured, the gratings with optimal spatial and temporal frequency for that unit were used and were varied over orientation. Each grating alternated between an amplitude of ±3 on a gray (0) background. Some units, especially in higher stacks, had weak or no responses to drifting sinusoidal gratings. To account for this, we excluded any units with a mean response (over time) of <1% of the maximum mean response of all the units in that stack. As a result of this, 0/100, 88/200 and 261/400 units were excluded from the 2^nd^, 3^rd^ and 4^th^ stacks, respectively.

We measured several aspects of the V1 neuron and model unit responses to the drifting gratings. For each unit, we measured the circular variance, orientation bandwidth, modulation ratio and direction selectivity.

As a control, we examined the RFs and responses to drifting gratings of each unit with its immediate input weights shuffled. In this case, the receptive fields lacked discernable structure, with only patchy spatial frequency and orientation tuning in response to gratings. There were very few orientation tuned (circular variance < 0.9) units with modulation ratios <1.

### Circular variance

Circular variance (CV) is a global measure of orientation selectivity. For a unit with mean response over time r_q_ to a grating with angle θ_q_, with angles θ spanning the range of 0 to 360° in equally spaced intervals of 5° and measured in radians, the circular variance is defined as [28]:

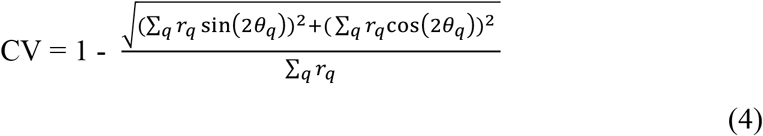

### Orientation bandwidth

We also measured the orientation bandwidth [28], which provides a more local measure of orientation selectivity. First, we smoothed the direction tuning curve with a Hanning window filter with a half-width at half-height of 13.5°. We then determined the peak of the orientation tuning curve. The orientation angles closest to the peak for which the response was 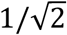 (or 70.7%) of the peak response were measured. The orientation bandwidth was defined as half of the difference between these two angles. We limited the maximum orientation bandwidth to 180°.

### Modulation ratio

We measured the modulation ratio of each unit’s response to its optimal sinusoidal grating. The modulation ratio is defined as:

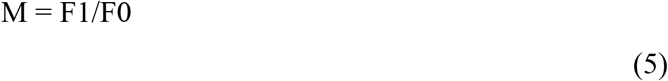

where F1 is the amplitude of the best-fitting sinusoid to the unit’s response over time to the drifting grating. F0 is the mean response to the grating over time.

### Direction selectivity index

To measure the direction selectivity index, we obtained each unit’s direction tuning curve at its optimal spatial and temporal frequency. We measured the peak of the direction tuning curve, indicating the unit’s response to gratings presented in the preferred direction (r_p_) as well as the response to the grating presented in the opposite (non-preferred) direction (r_np_). The direction selectivity index is then defined as:

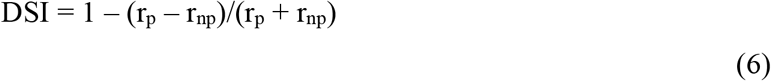

### Measuring end-stopping

In order to investigate end-stopping, we measured the responses of the hidden units to the same set of drifting gratings but with a circular mask applied to the inputs (e.g. Fig 5a, ii). Masks with a range of spatial extents were tested and the response of the units as a function of this spatial extent was measured (Fig. 5b).

### Sparse noise stimuli and two-bar maps

We measured the responses of the hidden units to ‘sparse noise’ (moving two-bar) stimuli [30]. Each stimulus contained a single oriented bar over the two most recent time-steps and a blank stimulus in the preceding time-steps. For each unit, the bar was oriented in the preferred orientation (as measured using drifting sinusoidal gratings) of the unit being probed. The length and width of the bar were limited to 50% and 10% of the unit’s spatial extent, respectively. This was typically enough for the bar to be longer than the unit’s spatial RF. In the first time-step with a bar, the center position of the bar (its x and y coordinate) was selected from a dense grid of spatial positions starting from the center of visual space and covering 1/3 of the unit’s spatial extent in each direction. In the second time-step, another bar position was selected from the same grid. This way, displacement of the bar from each grid position to each other grid position was used to stimulate the unit. To generate two-bar response maps, we measured the response of the unit as a function of the vertical and horizontal displacement (the starting position minus the end position) and then averaged over starting position. This gives a map of the unit’s response as a function of the displacement of the stimulus regardless of the starting position. We performed this procedure using all combinations of pairs of white (amplitude +3) and black (amplitude −3) bars on a gray (amplitude 0) background. This yielded four maps (white-to-white, black-to-black, white-to-black and black-to-white). We then summed the same contrast maps (white-to-white and black-to-black) and subtracted the opposite contrast maps (white-to-black and black-to-white) to yield the final two-bar map for each unit. This preserves directional responses while eliminating the responses that depend only on the spatial position of the bars in each frame [30].

Examining the two-bar maps, the position (0,0) indicates that the bar was in the same position in two successive frames, while the vertical and horizontal axes represent movement in these directions. Positive activity means that the unit was excited by movement in that direction, while negative activity indicates inhibition of the unit to movement in the given direction. A non-end-stopped unit will respond to any movement with a component in the preferred direction of the cell. This results in an elongated response profile on the two-bar map (Fig. 5c, right). An end-stopped unit will only respond to movement in the cell’s preferred direction, resulting in a two-bar map whose excitatory activity is limited to a more circumscribed region [30] (Fig. 5c, left).

### Drifting plaid stimuli

In order to test whether units were pattern selective, we measured their responses to drifting plaid stimuli. Each plaid stimulus was composed of two superimposed half-intensity (amplitude 1.5) sinusoidal gratings with different orientations. The net direction of the plaid movement lies midway between these two orientations (Fig. 6a). As with the sinusoidal inputs, plaids with a variety of orientations, spatial frequencies, temporal frequencies and spatial extents (as defined by the extent of a circular mask) were tested. For each unit, the direction tuning curves of the optimal plaid stimulus (that giving rise to the largest mean response over time) were measured (Fig. 6b,c).

### Code and data availability

All custom code used in this study was implemented in Python. We will upload the code to a public Github repository upon acceptance. The movies used for training the models are all publicly available at the websites detailed in the Methods. The V1 data used for comparison is available at http://www.ringachlab.net/lab/Data.html [28].

## Acknowledgements

Yosef Singer was supported by the University of Oxford Clarendon Fund, the Oppenheimer Memorial Trust and the Goodger and Schorstein Research Scholarship in Medical Sciences. Andrew King and Ben Willmore were supported by the Wellcome Trust (WT076508AIA, WT108369/Z/2015/Z). Nicol Harper was supported by a Sir Henry Wellcome Postdoctoral Fellowship (WT082692) and other Wellcome Trust funding (WT076508AIA, WT108369/Z/2015/Z), by the Department of Physiology, Anatomy and Genetics at the University of Oxford, by Action on Hearing Loss (PA07), and by the Biotechnology and Biological Sciences Research Council (BB/H008608/1).

## Author contributions

Conceptualization, N.S.H.; Methodology, Y.S., B.D.B.W., N.S.H; Investigation, Y.S; Visualization, Y.S.; Software, Y.S.; Writing – Original draft, Y.S.; Writing – Review & Editing, Y.S., B.D.B.W., A.J.K., N.S.H; Resources, A.J.K; Funding Acquisition, A.J.K; Supervision, B.D.B.W., A.J.K, N.S.H.

## Supplemental figures

**Figure S1.**
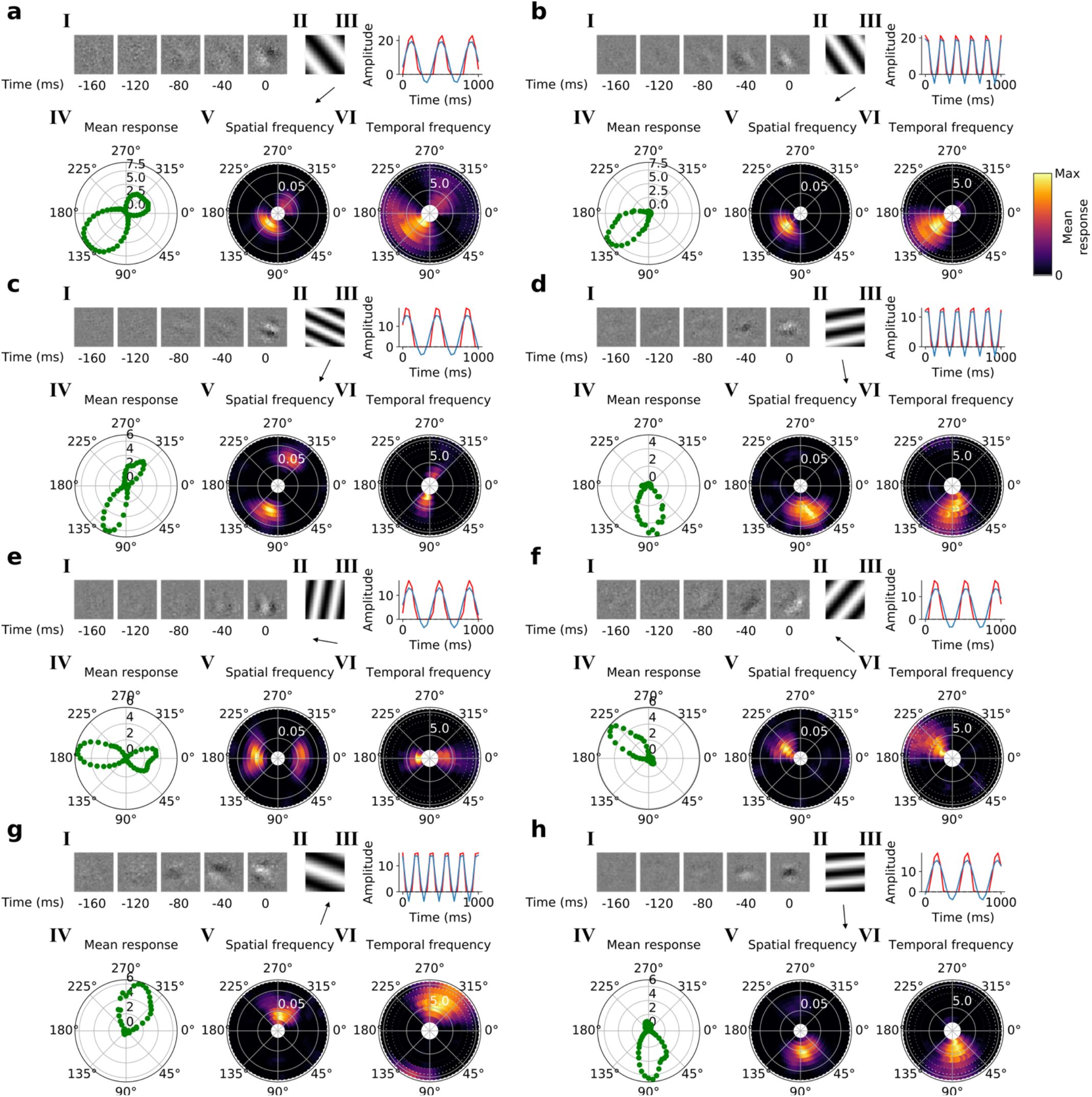
**Tuning properties of example units in stack 2. *a-h, I-IV*** as in Fig. 3*a-d*.

**Figure S2.**
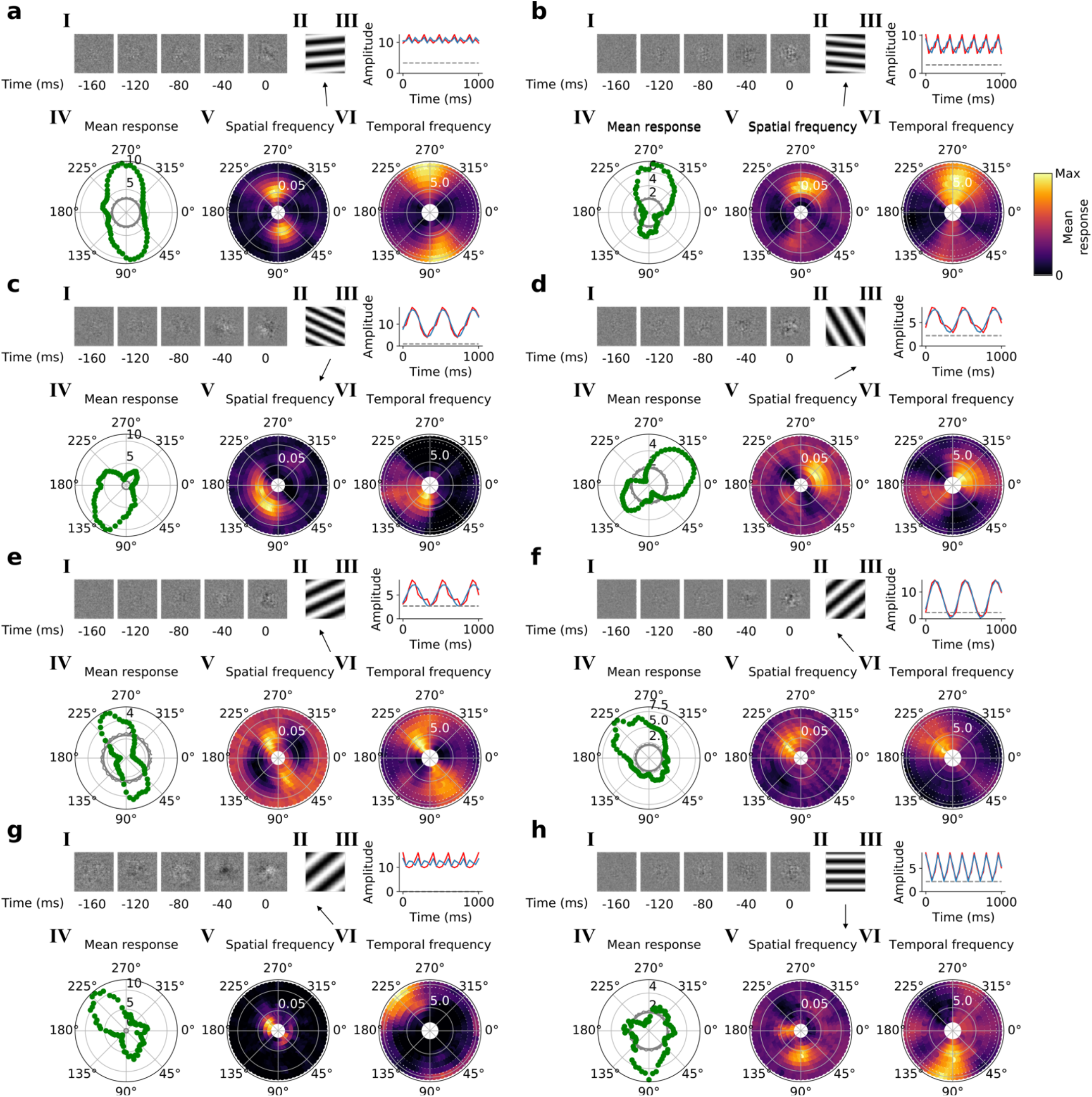
**Tuning properties of example units in stack 3. *a-h, I-IV*** as in Fig. 3*a-d*.

**Figure S3.**
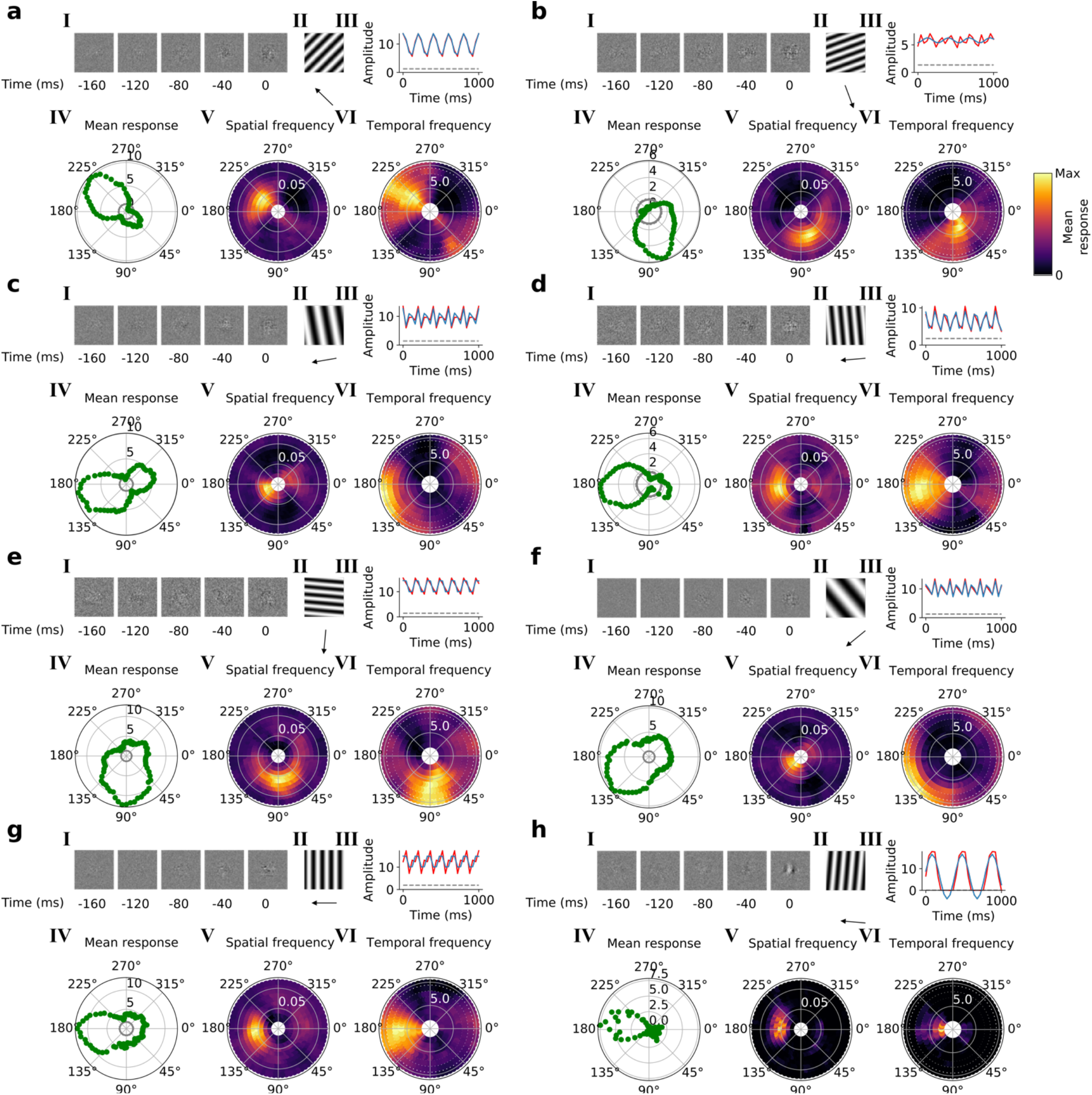
**Tuning properties of example units in stack 4. *a-h, I-IV*** as in Fig. 3*a-d*.

